# Spontaneous dynamics of hippocampal place fields in a model of combinatorial competition among stable inputs

**DOI:** 10.1101/2023.09.04.556254

**Authors:** Francesco Savelli

## Abstract

We present computer simulations illustrating how the plastic integration of spatially stable inputs could contribute to the dynamic character of hippocampal spatial representations. In novel environments of slightly larger size than typical apparatus, the emergence of well-defined place fields in real place cells seems to rely on inputs from normally functioning grid cells. Theoretically, the grid-to-place transformation is possible if a place cell is able to respond selectively to a combination of suitably aligned grids. We previously identified the functional characteristics that allow a synaptic plasticity rule to accomplish this selection by synaptic competition during rat foraging behavior. Here, we show that the synaptic competition can outlast the formation of place fields, contributing to their spatial reorganization over time, when the model is run in larger environments and the topographical/modular organization of grid inputs is taken into account. Co-simulated cells that differ only by their randomly assigned grid inputs display different degrees and kinds of spatial reorganization—ranging from place-field remapping to more subtle in-field changes or lapses in firing. The model predicts a greater number of place fields and propensity for remapping in place cells recorded from more septal regions of the hippocampus and/or in larger environments, motivating future experimental standardization across studies and animal models. In sum, spontaneous remapping could arise from rapid synaptic learning involving inputs that are functionally homogeneous, spatially stable, and minimally stochastic.

**Significance Statement:** In both AI and theoretical neuroscience, learning systems often rely on the asymptotic convergence of slow-acting learning rules applied to input spaces that are presumed to be sampled repeatedly, for example over developmental timescales. Place cells of the hippocampus testify to a neural system capable of rapidly encoding cognitive variables—such as the animal’s position in space—from limited experience. These internal representations undergo “spontaneous” changes over time, spurring much interest in their cognitive significance and underlying mechanisms. We investigate a model suggesting that some of these changes could be a tradeoff of rapid learning.

## Introduction

Hippocampal place cells differ functionally from upstream entorhinal grid cells beyond the lack of grid-patterned firing in space. Grid cells immediately lay out their characteristically regular firing pattern during the exploration of a novel environment (Hafting et al. 2005), whereas place cells can express their firing locations (“place fields”) more idiosyncratically (Wilson and McNaughton 1993; Frank, Stanley, and Brown 2004; Frank, Brown, and Stanley 2006; Monaco et al. 2014; Kim, Jung, and Royer 2020). Grid cells belonging to the same or different modules exhibit a coordinated switch of reference frame for spatial firing in response to contextual and perceptual changes (Stensola et al. 2012; Marozzi et al. 2015; Savelli, Luck, and Knierim 2017). In contrast, place cells can react independently to similar manipulations, collectively exhibiting “complex”, “partial”, or “global” place field remapping (Bostock, Muller, and Kubie 1991; Muller and Kubie 1987; Tanila, Shapiro, and Eichenbaum 1997; Skaggs and McNaughton 1998; Knierim 2002; Brown and Skaggs 2002; Leutgeb et al. 2005; Wills et al. 2005). The spontaneous remapping of place fields in the absence of experimental manipulations has also been increasingly documented, spurring speculation on the possible cognitive significance of the instability—or flexibility— of the place cell code (Ludvig 1999; Manns, Howard, and Eichenbaum 2007; Mankin et al. 2012; 2015; Rubin et al. 2015; Hainmueller and Bartos 2018; Muzzio 2018; Khatib et al. 2023). Non-spatial inputs or spatial inputs other than grid cells could be responsible for some of these functional differences between grid and place cells (Muzzio et al. 2009; Kentros et al. 2004; Tsao et al. 2018; Diehl et al. 2018; Save, Nerad, and Poucet 2000; Fischler-Ruiz et al. 2021). In this study, we explore the counterintuitive idea that spatially stable grid cells, too, could contribute to the dynamic character of hippocampal place fields as a consequence of the rapid learning required for their generation on a behaviorally realistic timescale.

Different lines of evidence have indirectly implicated the Medial Entorhinal Cortex (MEC) and its grid cells in the formation of hippocampal place fields. Although inactivation of the medial septum has been shown to spare place fields despite causing a severe disruption of grid firing patterns (Brandon et al. 2014; 2011; Koenig et al. 2011), it prevents the emergence of place fields for at least three days in a larger, novel environment (Wang et al. 2015). Similarly, MEC lesions dramatically affect place fields in a novel environment, but not in a slightly smaller and familiar environment (Hales et al. 2014). Developmentally, when the grid-cell system has not yet reached functional maturation, place fields in the center of the apparatus are found to be more unstable than peripheral ones (Muessig et al. 2015). Moreover, genetic manipulations of grid cells’ spatial scales cause analogous changes in centrally located place fields (Mallory et al. 2018), and chemogenetic manipulations of firing rate levels in MEC can cause place cells to remap (Kanter et al. 2017). Together, these observations point to a major influence of grid cell inputs on hippocampal place fields, which is obscured in small environments that scarcely accommodate grid patterns (see, for example, Quirk et al. 1992; Fyhn et al. 2004; Hargreaves et al. 2005 vs. Hafting et al. 2005, and further discussion in Savelli and Knierim 2019.)

Theoretically, the nonlinear sum of suitably aligned grids can yield a peak of activity where their vertices maximally overlap (Solstad, Moser, and Einevoll 2006). We previously showed how a model place cell can generate a place field by becoming selectively responsive to one among multiple possible alignments of its grid inputs via synaptic plasticity (Savelli and Knierim 2010). Here, we investigate the possibility that the same learning dynamics could account for spontaneous remapping or drift of established place fields through continued synaptic competition among grid inputs.

## Essential features of the model

The model and simulations presented here differ from those described before (Savelli and Knierim 2010) mainly in two aspects: the use of additional environments of varying sizes and the modular/topographical organization of grid-cell inputs to place cells.

Because environment size appears to be a crucial factor in how grid cells influence place fields, we ran the model on trajectory data obtained from open-space arenas of varying sizes. On the smaller end, we reused the same trajectory from the previous study in a 60x60cm box (“Box60”). The new environments are a 68x68cm box (“Box68”), a 76cm-diameter walled cylinder (“Cyl76”, with area nominally equal to Box68—these two types of arenas have been conventional in many classic place cells studies), a 100x100cm platform (“Plat100”), and a 137x137cm platform (“Plat137”). The platforms have no walls, marginally extending the area potentially sampled by the rat (Monaco et al. 2014).

The pool of grid cells providing input to the model was segregated into seven “modules” (200 grid cells per module) postulated to be located in progressively more ventral locations of the Medial Entorhinal Cortex (MEC) (Fig. 1A). The scale increased discretely from one module to the next according to a geometric progression (inter-vertex spacing ranging from 40cm to 328cm, progression ratio 1.42), based on extrapolations suggested by experimental observations (Stensola et al. 2012). Grid cells in the same module shared the same orientation and scale. Module orientations did not appear to influence the results of this study (not shown). We hence report results based on a choice of values consistent with the documented tendency of grids to slightly misalign relative to the cardinal axes of a symmetric environment (Stensola et al. 2015) (Fig. 1B, left). The individual phases of grids from the same module were drawn from a uniform distribution over a hexagonally shaped interval representing the first 2D period of the grid (Fig. 1B, left). Any inhomogeneity in the spatial density of these samples was inevitably repeated in environments capable of accommodating multiple grid periods (Fig. 1B, right). The momentary animal location modulated the instantaneous firing rate of each grid cell’s spike train, peaking at 20Hz in the center of each vertex. Grid inputs thus provided spatially stable patterns of excitation to simulated place cells, with the only noise factor provided by the spike jitter inherent in the statistical (Poisson) generation of spike timings.

**Figure 1.**
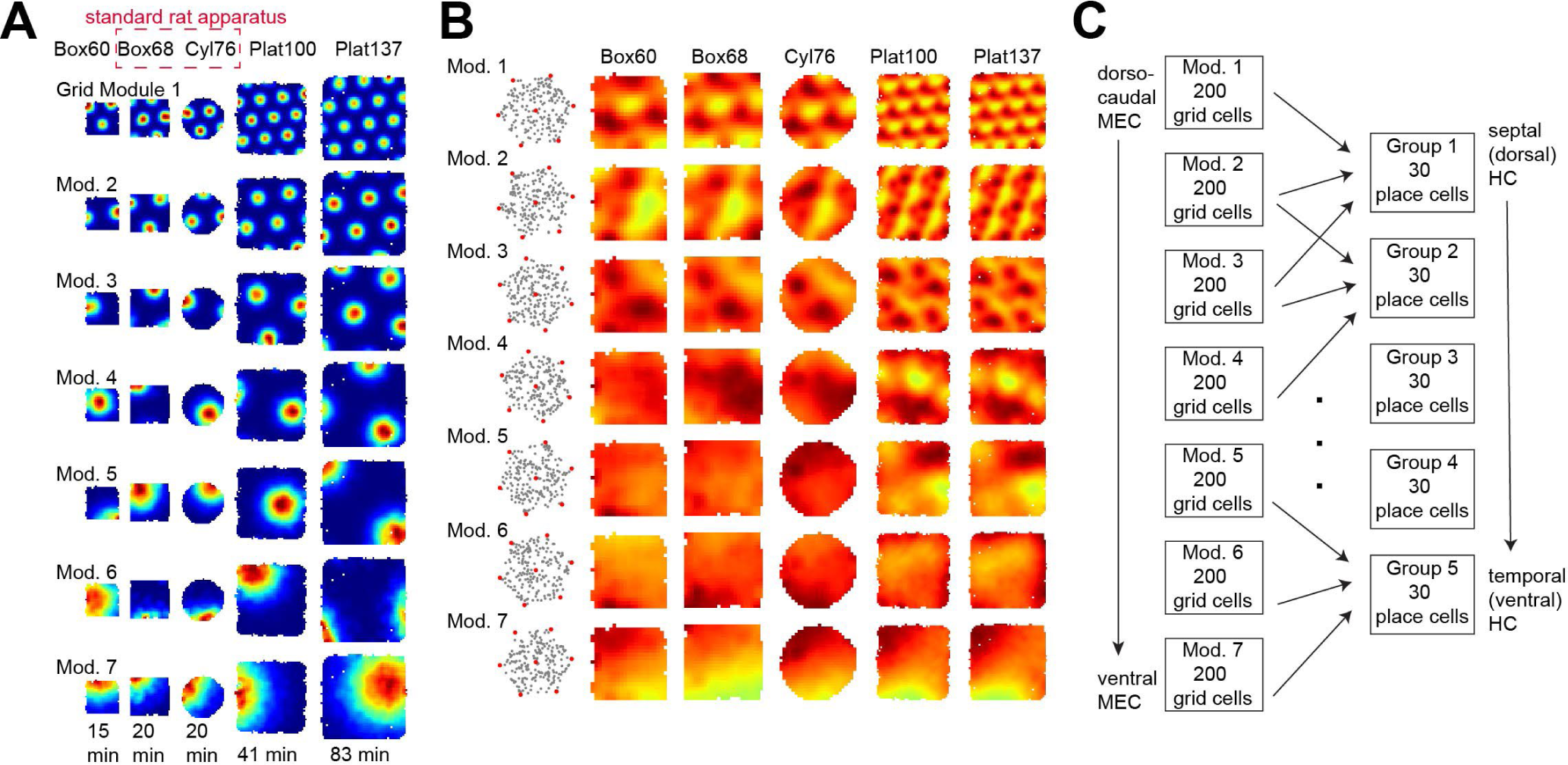
Model inputs. **(A)** An example of a grid cell rate map for every module-environment combination. More temporal modules have larger grid scales. The duration of each simulated session was proportional to the area of the environment. **(B)** (Left) Shared orientation and phases of all grids simulated in each module. Phases are represented by one dot within one hexagonal grid period, which is tilted along the orientation shared by all grids of the module. The phases are drawn from a uniform distribution and inhomogeneities in their spatial density arise by chance. (Modules’ period sizes have been equalized to aid visibility.) (Right) Sum of all the grids from each module plotted in each environment. (Environments’ sizes have been equalized here to aid visibility, but grid scales are relative to the environment.) Note how the inhomogeneities recur for as many grid cycles as the environment can accommodate. **(C)** Schematic of the modular/topographical organization of the grid-cell inputs to place cells. Each group receives input from three consecutive modules.

The input to each simulated hippocampal cell was restricted to 60 projections from grid cells randomly chosen from three consecutive grid modules (Fig. 1C). This choice was motivated by the topographical organization of MEC-HC projections (Fyhn et al. 2004), the documented anatomical contiguity and degree of overlap among modules (Stensola et al. 2012), and by reports of up to three different grid scales being observed simultaneously at a single recording site in MEC (Savelli, Luck, and Knierim 2017). Consequently, we simulated 5 groups of place cells intended to represent populations of place cells topographically arranged along the septo-temporal axis of the hippocampus (30 place cells per group, Fig. 1C). Place cells were modeled as integrate-and-fire spiking units with synaptic conductances (one conductance per grid cell synapsing on the place cell). At each input spike occurrence, the synaptic conductance transiently increases by a variable amount that plays the role of “synaptic weight” and is subject to plasticity.

Plasticity follows a “postsynaptically-gated” synaptic plasticity rule:

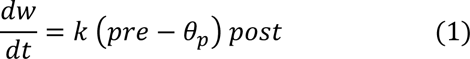

where *w* is the weight, *pre* is the presynaptic firing rate, *post* is the postsynaptic firing rate, *k* is the learning rate factor, and *θ_p_* is a threshold on the presynaptic firing rate (5 Hz). Postsynaptic activity is required for the right-hand side of the differential equation to be different from 0 and trigger synaptic modification. The direction of change is determined by the presynaptic firing rate: if *pre* is greater than *θ_p_*, the right-hand side of the equation is positive and the synapse is potentiated; otherwise, the synapse is depressed. As we showed before (Savelli and Knierim 2010), the rate of learning *k* must be sufficiently high to enable to formation of place fields during a typical behavioral session. However, the firing rate traces *pre* and *post* were computed by convolving the presynaptic and postsynaptic spike trains with an exponential kernel with time constant set to 100 ms (i.e., approximating calcium dynamics).

## Place fields emerge from diffuse input drive in all environments

The sum of all grid inputs randomly assigned to a place cell amounts to a non-uniform but diffuse landscape of excitatory drive regardless of environments (Fig. 2, “input drive” maps). Restricting the place cell’s firing to a well-defined place field requires isolating a combination of grids whose vertices are suitably aligned to produce a spot of maximum excitation (Solstad, Moser, and Einevoll 2006; Franzius, Vollgraf, and Wiskott 2007; Blair, Welday, and Zhang 2007; Azizi, Schieferstein, and Cheng 2014) (Fig. 2, “rate” maps). Synaptic plasticity can select for such a combination during an animal’s exploration of an environment by rapidly potentiating its input grids while depressing all the other (non-aligned) inputs (Savelli and Knierim 2010). The synaptic rule (Eq. 1) successfully accomplishes this task to produce place fields in all of the new environments (Fig. 2A) with the discrete/topographical organization of the MEC-HC forward projections (Fig. 1C) considered here. In most cases, this process refines and amplifies one or more of the peaks in the corresponding input drive map (compare input drive vs. rate maps in Fig. 2A). In other cases, place fields emerge in less obvious regions relative to the input drive map (Fig. 2B).

**Figure 2.**
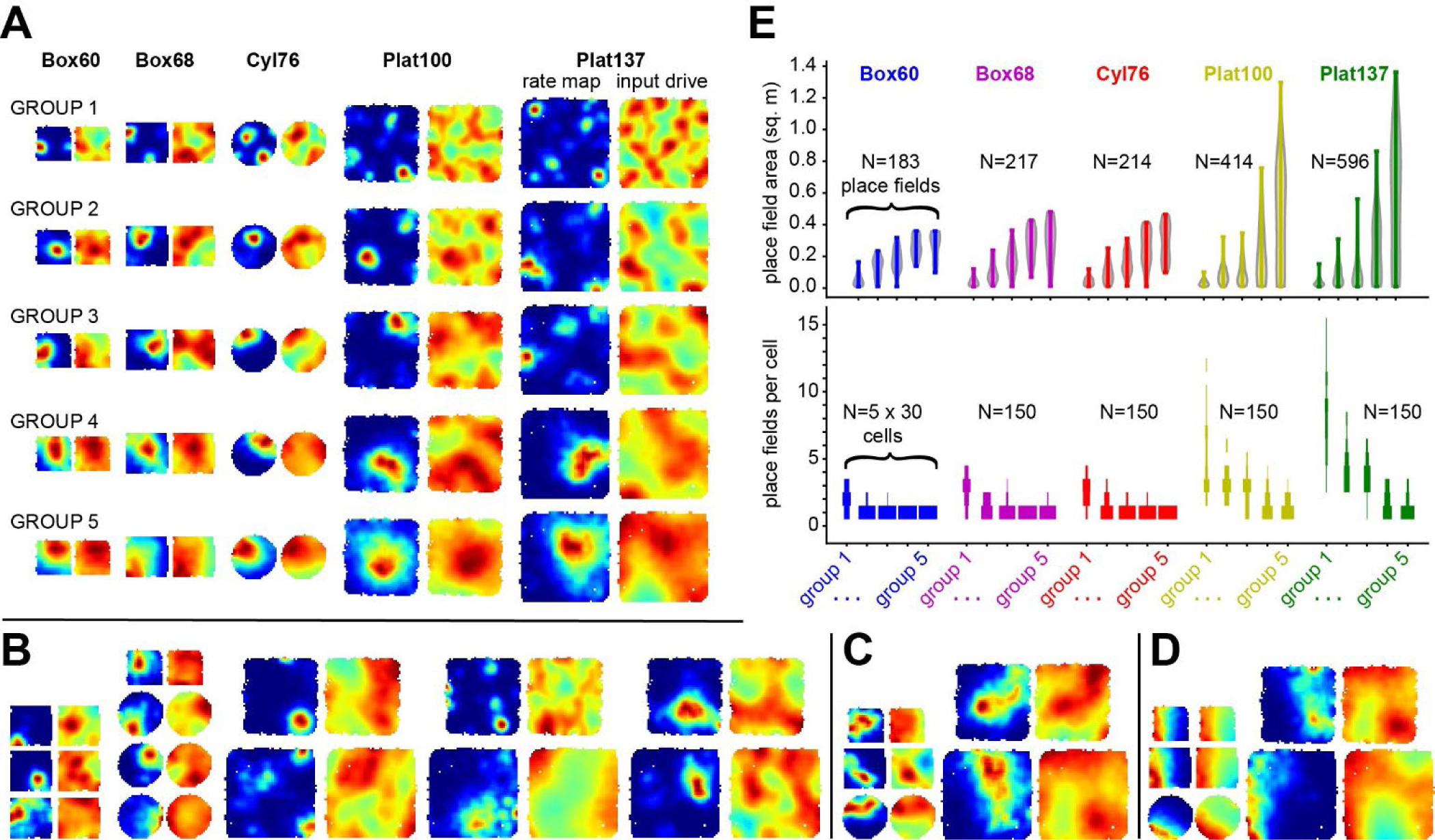
Place cells generated by the model. **(A)** An example of a place cell for every group-environment combination. For each place cell a firing rate map (left) and input drive map (right) is given (individually autoscaled). The input drive map is the sum of all grid input to the cell. **(B)** Examples of place cells whose firing rate patterns would be difficult to predict from their input drive maps. **(C)** Examples of place fields with irregular shapes. **(D)** Examples of place cells firing along the boundaries of the environment. **(E)** (Top) Place field sizes for all place fields from all co-simulated place cells in each environment by group. Violin plots represent distribution densities. Temporal groups express larger place fields, reminiscent of the septo-temporal topographical organization found in the real hippocampus. (Bottom) Number of place fields per cell from all co-simulated place cells in each environment by group. Each discrete violin plot is a histogram representing cell counts in each horizontal bin. Septal groups express more fields per cell.

Place fields with (roughly) round outlines or with more irregular shapes are generated across different place cells that are co-simulated in the same environment (Fig. 2A vs 2C). Some place fields were reminiscent of boundary cells (Fig. 2D) (Savelli, Yoganarasimha, and Knierim 2008; Solstad et al. 2008; Lever et al. 2009), even though only grid inputs were used. All of the maps for all cells simulated in each environment are reported in SFig2_1-5. Peak and average firing rates across all 750 cells were physiologically plausible (peak firing rate: mean 12.2 Hz, std 5.8 Hz; average firing rate: mean 2.6 Hz, std 2.6 Hz).

## Place field size and numerosity are topographically organized

We detected place fields as regions of elevated firing on rate maps (as conventional in the experimental literature, see method details) and quantified their properties. At least 70% of firing occurs within the place fields in 648 out of the 750 cells simulated across all environments (mean 82.5%, std 12%), demonstrating that these fields account for most of the simulated cells’ activity. The range of place field sizes is larger in more temporal groups and in larger environments that allow for the expression of the largest fields (Fig. 2E Top). This “topographical” organization of place field size is consistent with experimental observations (Jung, Wiener, and McNaughton 1994; Kjelstrup et al. 2008) and results from the organization we imposed on grid cells’ scale and projections to place cells. Interestingly, the number of place fields per cell is similarly topographically organized, with cells in septal groups generating more fields (Fig. 2E Bottom). For instance, place cells in groups 1 and 2 generate multiple, very small place fields in all environments (Fig. 2A, SFig2_1-2). In some environments, multiple fields are arranged in quasi-regular patterns that are highly reminiscent of grid cell firing patterns (Plat100 and Plat137, SFig2_4,5). This quasi-periodic structure is already evident in the input drive map, reflecting periodic inhomogeneities of the summation of grids assigned to the cell (Fig. 1B), rather than a dominant input from a single grid emerging from the synaptic learning process. (The regular firing pattern can be highly unstable throughout the session, reflecting competition among different grid inputs, see next sections.) These uncharacteristic place cells are predicted to be located at the most septal end of the hippocampus, an area that to the best of our knowledge has not been targeted by experimental studies employing open arenas at least as large as Plat100.

Place cells in groups 3 and 4 express typical place fields in typical experimental environments (Box68, Cyl76, Fig. 2A, SFig2_2,3) but often produce multiple place fields in larger environments (Plat100, Plat137, Fig. 2A, SFig2_4,5). In experimental studies that recorded place cells from rats running in environments similar to Plat137 in size (Fenton et al. 2008; Park, Dvorak, and Fenton 2011; Wang et al. 2015), place fields appeared comparable to those from groups 2-4.

## Competing grid combinations can account for place field instability of different kinds and degrees

Newly formed place fields could be destabilized by continuing synaptic competition. If grid inputs other than those driving the place field retain sufficient synaptic efficacy and combine to drive out-of-field firing, they could trigger the formation of a new field and extinguish the original one. Assessing this possibility in our model requires analysis with more temporal resolution than averaging a cell’s spatial activity throughout the session into a single firing rate map. One common approach has been to compute two rate maps corresponding to the first and second halves of the recording session. These two partial rate maps are then compared visually or quantitatively by a bin-wise Pearson correlation (“rate map stability score”, RMS, ranging from -1 to 1). This *cell-based analysis* provides a rough estimate of the stability of the cell’s activation in space, but it depends on an arbitrary timepoint and generally cannot provide a clear timeline of the development and stability of the cell’s individual place fields. We therefore developed a *field-based analysis* by computing (i) a temporal trace of the cell’s firing rate within a place field (activity trace) and (ii) an analogous trace of the rate of animal’s presence inside the same region (occupancy trace). Visual comparison of these two traces reveals when a place field first emerged and whether/when its activity stopped/resumed, while taking into account when the animal’s location gave the place cell the opportunity to fire in the field. We quantified the temporal alignment of the fluctuations of the activity vs. occupancy traces by computing the normalized dot product of the two traces (“place field stability score”, PFS, ranging from 0 to 1).

A clear example of place field remapping in Box68 captured by both analyses is illustrated in Fig. 3A. Place field #1 (blue trace) emerges from diffuse input drive within the first minute of exploration as the initially uniform synaptic weights split into potentiated and depressed. The field and weights remain stable until about midsession when this field becomes inactive and place field #2 (red trace) first emerges in a different location. Both regions were well sampled by the rat before and after this transition as shown by their respective occupancy traces. The cell thus undergoes spontaneous, abrupt remapping. The transition point is characterized by a dramatic redistribution of synaptic weights across the inputs, which reshapes the landscape of synaptic drive (see “instantaneous weighted sums of input grids” running on top of Fig. 3A). The new place field remains largely stable for the remainder of the session, although minor synaptic reorganization events affect its exact shape. Note how these events do not always affect all the inputs at once, as some inputs remain potentiated. We quantified synaptic changes in time by correlating the synaptic weight vectors at a 10s lag (Pearson correlation, illustrated below the temporal plot of synaptic weights vector in Fig. 3A).

**Figure 3.**
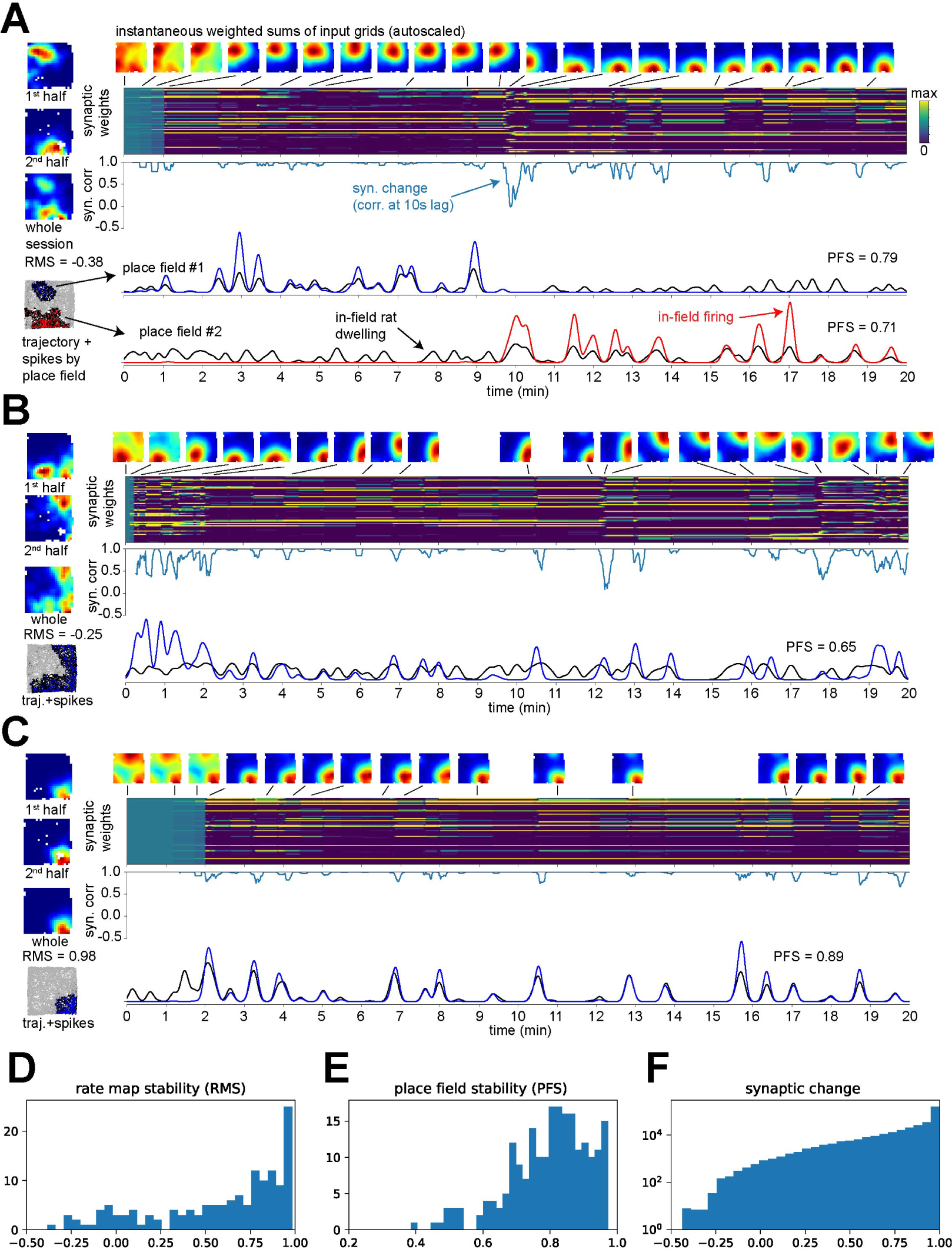
Place field stability in Box68. **(A)** Example of place cell from group 3 that remaps midsession. (Left) Firing rate maps for the first half, second half, and whole session (red: max firing, blue: no firing; color code autoscaled individually in each map). Bottommost is a spatial raster plot of the trajectory and spikes (light grey), but the spikes assigned to the first identified place field are in blue and those assigned to the second place field are in red; trajectory segments occurring within these place fields are in black. (Right) Color-coded evolution of the synaptic weight vector for the 60 grid-cell inputs. Weighted sums of these inputs illustrate synaptic drive as a function of space at salient timepoints based on instantaneous readings of the synaptic weight vector (minimaps are individually autoscaled). Synaptic change is traced over time as correlations between pairs of synaptic weight vectors 10s apart taken every 500ms. Two bottommost plots illustrate activity traces in color and occupancy traces in black for both place fields (see main text). Note extinction of the first place field and emergence of the second coincide. **(B)** Example of drifting place field from a group 3 place cell, illustrated as in A. **(C)** Example of very stable place field from a group 3 place cell, illustrated as in A. **(D)** Distribution of rate map stability scores for all cells co-simulated in Box68. **(E)** Distribution of place field stability scores for all identified place fields from all cells co-simulated in Box68. **(F)** Distribution of 10s-lag synaptic vector correlations taken every 500ms from all cells co-simulated in Box68, note logarithmic scale.

In another example (Fig. 3B), after the initial place field is formed, synaptic changes cause it to drift in space. It appears to “crawl” along the walls. A third place cell (Fig. 3C) generates a single and very stable place field. Its synaptic weights undergo negligible perturbations after its formation. These three examples are all from group 3 of the same simulation in Box68. These cells therefore only differed in the inputs assigned to them randomly as all other parameters remained the same. In particular, their qualitatively different responses developed along the same animal trajectory. In Fig. 3D,E, the stability of all co-simulated place cells/fields in Box68 is quantified. Both RMS and PFS take values over a large range reflecting varying degrees of stability. The distribution of synaptic changes taken at regular timepoints from all cells is strongly skewed toward high correlation values (Fig. 3F, note logarithmic scale), reflecting synaptic stability most of the time for most of the cells. Still, the distribution covers the whole range of values stretching to negative numbers (all-encompassing synaptic changes), reflecting how synaptic reorganization of varying degrees can occur in a 10s timeframe. This quantification confirms that the combinations of grids controlling a place cell’s firing before and after a synaptic reorganization event are not necessarily fully disjoint.

The same variety of responses can be appreciated from similar examples and quantifications of co-simulated cells in the other environments (Fig. 4-6). For each environment we picked similar examples of remapping, drifting, or stable fields that convey further nuances in the dynamics of place fields. Figs. 4A and 5A show alternance between two fields, with multiple remapping events. Fig. 6A shows remapping involving 7-8 place fields. The place fields in Figs. 4B and 5B are distorted by alternate/jittery drift. The place field in Fig. 4C is quite stable but exhibits minor shifts and lapses in firing. Another field in Fig. 6B shows a ∼20-minute lapse in activity but is otherwise very stable.

**Figure 4.**
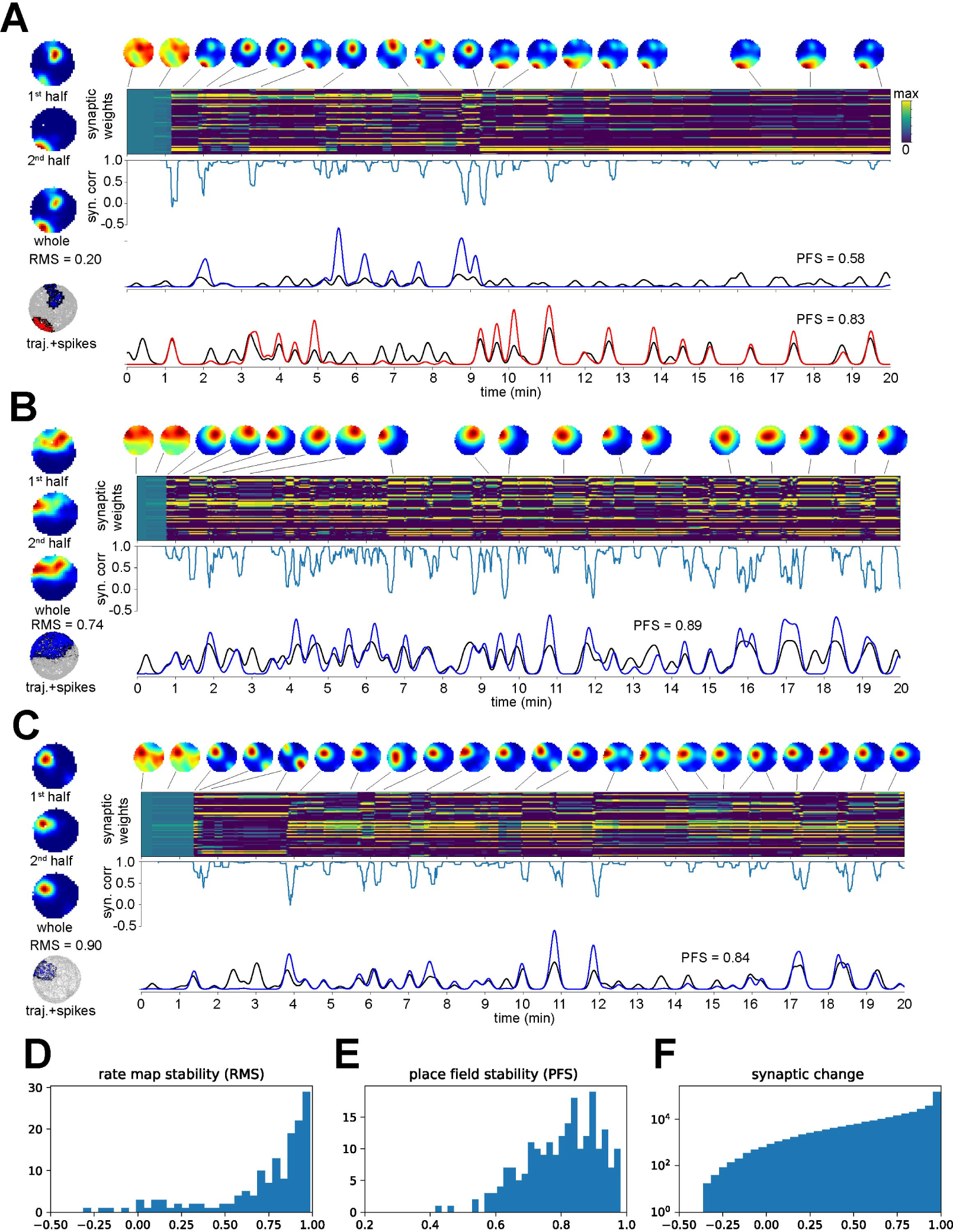
Place field stability in Cyl76. Examples and quantifications from the simulation in Cyl76 illustrated as in Fig. 3. The place cell in A (group 1) remaps more than once. The place field in B (group 2) drifts back and forth along an irregular path. The place field in C (group 1) is generally stable with short lapses in firing due to minor shifts of the place field caused by synaptic reorganizations.

**Figure 5.**
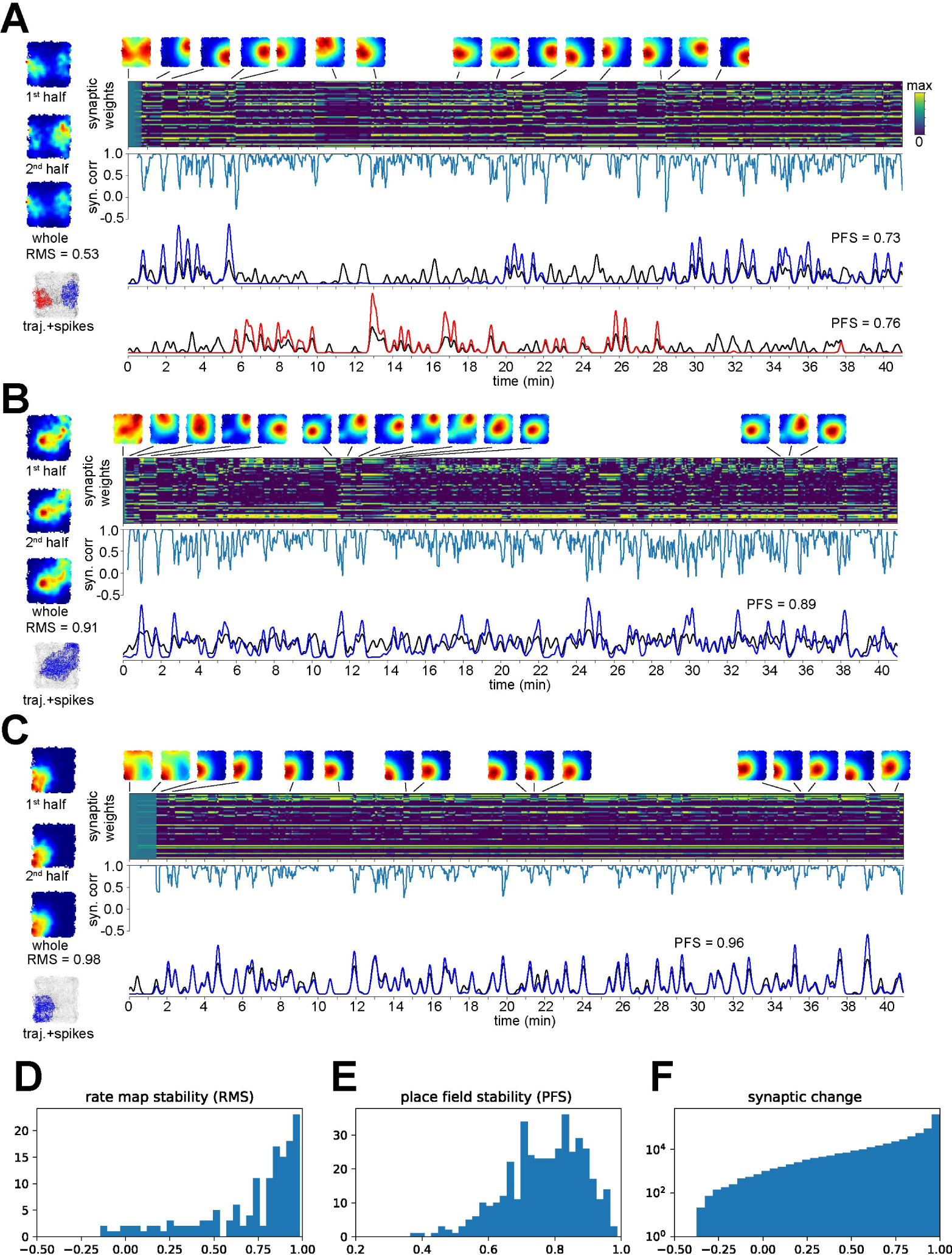
Place field stability in Plat100. Examples and quantifications from the simulation in Plat100 illustrated as in Fig. 3. The place cell in A (group 4) remaps more than once. The place field in B (group 3) drifts back and forth along an irregular path. The place field in C (group 4) is stable despite minor synaptic reorganizations.

**Figure 6.**
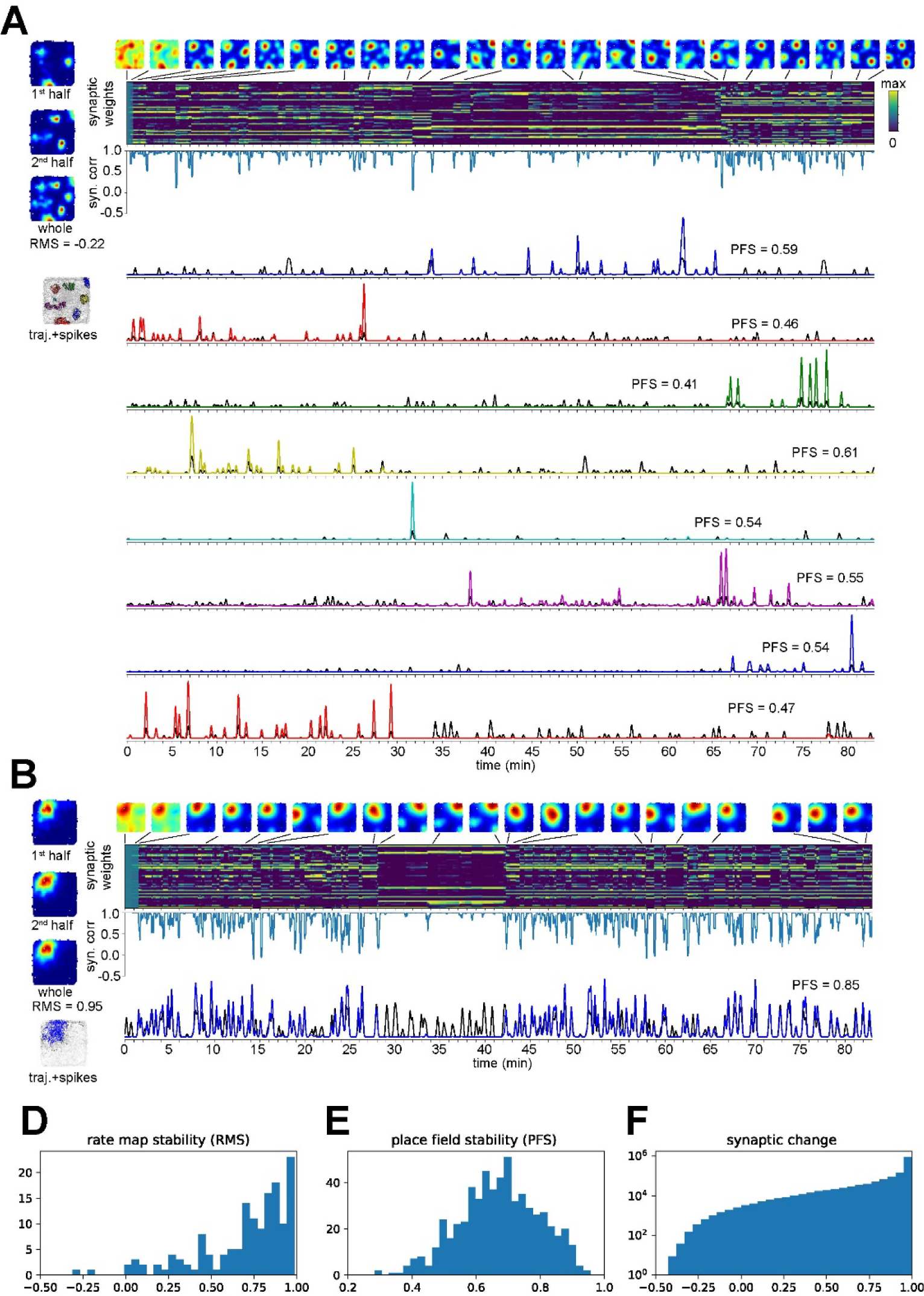
Place field stability in Plat137. Examples and quantifications from the simulation in Plat137 illustrated as in Fig. 3. The place cell in A (group 1) generates 7-8 place fields that are active during different but sometimes overlapping epochs of the session. The place field in B (group 3) is stable except for a long lapse in firing.

## Environment and place field size influence stability

Intuitively, larger environments and/or inputs from higher-resolution (smaller-scale) grid cells should offer more opportunities for spatial competition among different combinations of grids aligning in different locations. This intuition is confirmed by our quantifications of place field stability. Place fields produced by place cells from septal groups, which receive inputs from higher-resolution grid cells, are generally less stable than those from temporal groups (Fig. 7A). The same quantifications regrouped by environment show lower stability of place fields in larger environments (Fig. 7B). The longer durations of the sessions in the larger environments could contribute to this effect, although they allow for comparable spatial sampling and opportunities for remapping. We re-ran an extended simulation in Box68 and a shortened simulation in Plat137 so that both sessions were 40 minutes long. The stability distributions still diverge, albeit to a lesser extent (Fig. 7C).

**Figure 7.**
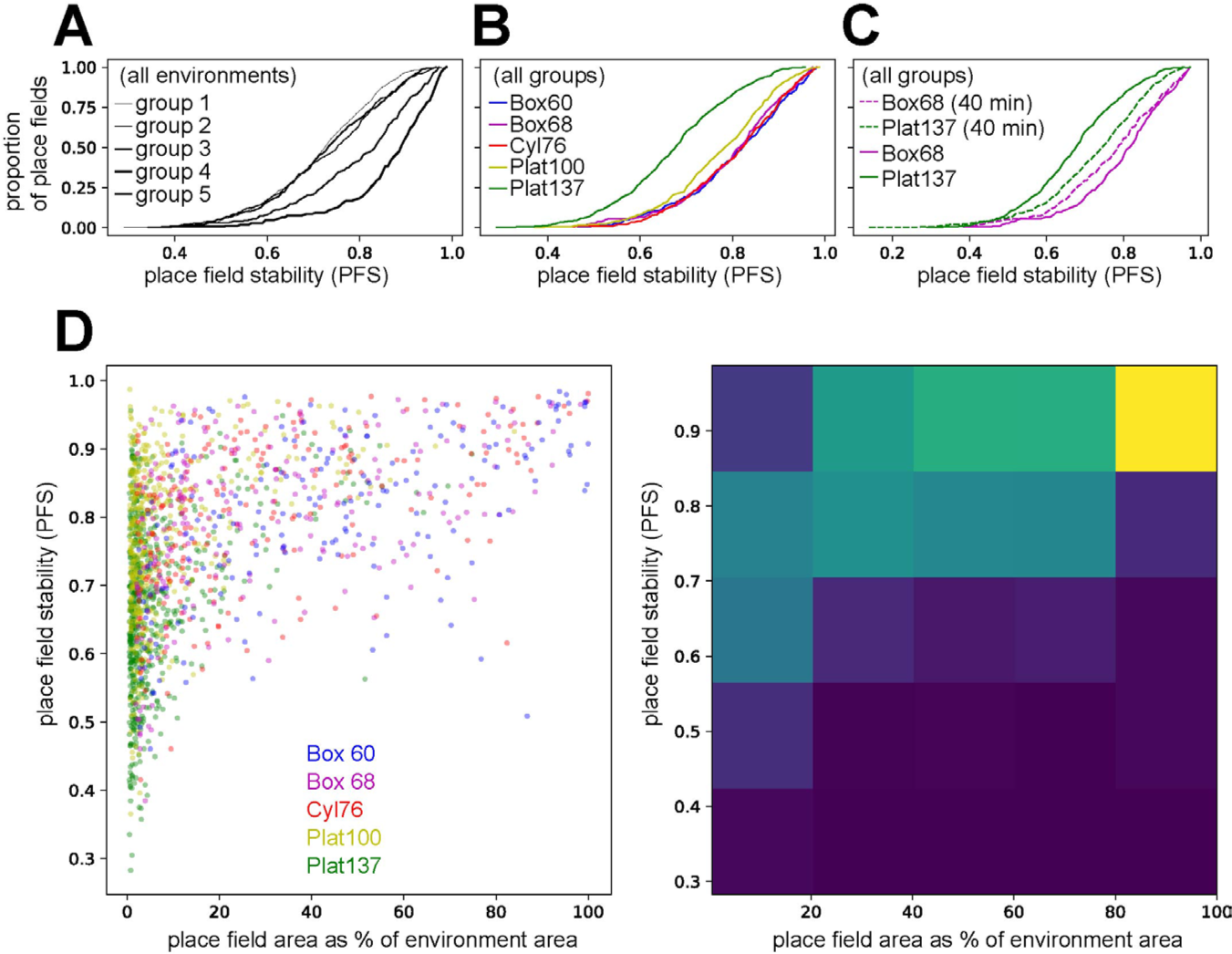
Spatial scale effect on stability. **(A)** PFS cumulative curves for all place fields from all simulations by place cell group. Note greater stability in the temporal groups. **(B)** PFS cumulative curves for all place fields from all simulations by environment. Note greater instability in the larger environments. **(C)** PFS cumulative curves for Plat137 and Box68 as in B plus a truncated version of Plat137 and an extended version of Box68 that have equal duration. **(D)** (Left) Each place field is colored by environment and plotted by its PFS and relative size. The most unstable fields are concentrated among those that are small relative to the environment, although not all such fields are necessarily unstable. (Right) The same data is plotted as a 2D color-coded histogram. The bins are normalized by column, to account for the decreasing availability of data points moving from left to right along the x -axis. Note the roughly “opposite upper triangular” structure illustrating the relationship between stability and relative place field size.

Since both environment size and septo-temporal location have an effect on place field stability, and place field size depends on septo-temporal location (Fig. 2E), we examined their combined effect by looking at the relationship between a place field’s stability and its size relative to the size of the environment (Fig. 7D). These plots show that unstable place fields are primarily found among those that are small relative to the environment. The model therefore predicts that differences in stability could be found along the septo-temporal axis of the hippocampus, with some place fields at its most septal end being especially unstable in environments larger than typical apparatus.

## Discussion

In our model, synaptic competition mediated by plasticity is presumed to tune place cells’ spatial activity from grid cell inputs during typical laboratory foraging behavior (Savelli and Knierim 2010). Here, we demonstrated that this competition can outlast the formation of well-defined place fields, leading to their spatial reorganization. Thus, even inputs that are functionally homogenous, spatially stable, and minimally stochastic can theoretically make place cells susceptible to spontaneous remapping when synaptic plasticity is active. Our simulated sessions lasted longer than most experiments in environments of comparable size, amounting to multiple typical experimental sessions spanning hours or days strung together. Because elapsed time per se had no active role in our model, our results are consistent with the role of continuing active experience in the drift of place-cell maps (Khatib et al. 2023). These computational insights do not rule out the possible involvement of additional mechanisms or inputs in the instability (or flexibility) of the hippocampal spatial representation (Olypher, Lánský, and Fenton 2002; Kentros et al. 2004; Muzzio et al. 2009; Tsao et al. 2018; Diehl et al. 2018; Low et al. 2022; Mau et al. 2018).

The synaptic reorganization underlying the dynamic character of our simulated place fields involve different—but not always altogether disjoint—combinations of grid inputs, each encompassing grids whose vertices coincide in at least one location. Different combinations of grids thus compete to control postsynaptic activation in space. As a result of this process, co-simulated place cells produce a mix of stable and unstable spatial correlates in the same environment, mimicking a variety of experimental observations. Highly stable place fields are consistent with the classic view of place cells as providers of a reliable position signal. However, minor variability or lapses in spatial firing in otherwise stable place fields are reminiscent of the overdispersion phenomenon (Fenton and Muller 1998; Olypher, Lánský, and Fenton 2002), whereby the variance of a place cell’s firing rate across multiple visits to the same location is found to be in excess of that of a Poisson spiking model (which we imposed on the grid inputs). Other place fields exhibited alternating or progressive drift between two nearby locations, giving rise to irregularly shaped place fields, which are not uncommon (e.g., Lever et al. 2002) and may have attracted less experimental attention than place fields conforming to stricter qualitative and quantitative spatial tuning criteria. When a similar dynamic involved locations farther apart, the place fields abruptly relocated, as has been observed in real place cells undergoing spontaneous remapping (Ludvig 1999; Mankin et al. 2012; 2015; Rubin et al. 2015; Khatib et al. 2023) or during a volatile process of initial formation (Wilson and McNaughton 1993; Frank, Stanley, and Brown 2004; Frank, Brown, and Stanley 2006). In our model, this response heterogeneity is entirely accounted for by the different sets of grid cells randomly assigned as inputs to co-simulated cells. No other biophysical parameter or behavioral input changed across these cells.

A specific prediction of our model concerns the interplay of the spatial scales of the environment vs. the environment’s representation by grid/place cells. More input competition—and therefore instability—is expected when the animal samples larger environments (or open-space arenas as opposed to tracks), or if place cells are recorded from more septal regions of the hippocampus receiving higher-resolution grid input. Both factors can multiply the locations at which grids can re-combine into different subsets with coinciding vertices. Indeed, our simulations predict an uncharacteristic proliferation of potentially unstable place fields in environments of tractable size (∼1.5mx1.5m) for place cells putatively located at the most septal end of hippocampus. To our knowledge, this anatomically small region has not been targeted by single-unit recordings (but see Maurer et al. 2006 for recordings using linear tracks that are more septal than in most studies). Similarly, in very large open-space environments, instability could affect the multi-field patterns characterizing cells recorded from more typical hippocampal locations (Saxena, Barde, and Deshmukh 2018; Harland et al. 2021; Tanni, de Cothi, and Barry 2022). Conceptually, the predicted topographical organization of place cell stability would be of interest for any functional interpretation of this property (Ludvig 1999; Manns, Howard, and Eichenbaum 2007; Mankin et al. 2012; Rubin et al. 2015; Hainmueller and Bartos 2018; Muzzio 2018; Khatib et al. 2023). Methodologically, this prediction motivates the standardization of both the environment size and recording locations along the hippocampus’ septotemporal axis in studies of place cell stability. Likewise, experimental comparisons involving different species (Rotenberg et al. 2000; Mou et al. 2018; Hok et al. 2016; Wirtshafter and Disterhoft 2022; Liberti et al. 2022; Payne, Lynch, and Aronov 2021; Agarwal et al. 2023) would require determining an appropriate rescaling of the recording environment calibrated to the range of spatial scales naturally expressed by the species’ grid and place cells. The modularity of grid size has been well characterized in the rat (Stensola et al. 2012; Savelli, Luck, and Knierim 2017), and less systematically in the mouse (Fyhn et al. 2008; Gardner et al. 2022), but not in other species.

The model is potentially relevant to the interpretation of other experimental observations. As we showed before, the early stage of exploration can bias where place fields are likely to appear, an effect mitigated by incorporating collateral inhibition in the model (Savelli and Knierim 2010). Similarly, uneven spatial sampling might favor combinations of grids active in oversampled locations. This might be relevant to place cell remapping following increased exploration of manipulated items (Fyhn et al. 2002; GoodSmith et al. 2022; Deshmukh and Knierim 2013), or to conflicting observations of place cells’ overrepresentation of goal locations across studies using different behavioral protocols (Hollup et al. 2001; Duvelle et al. 2019; Dupret et al. 2010). Moreover, sampling a new location could bring a new combination of grids into effective alignment, or extend the region of influence of an existing one by recruiting additional inputs, possibly triggering synaptic reorganization that leads to persistent modifications of the spatial response of the cell. This might be one factor contributing to the emergence of new place fields after increased off-track “scanning” by rats, in addition to covert attentional/perceptual processes (Monaco et al. 2014, see also Goble, Møller, and Thompson 2009).

Committing the physiological interpretation of the model to a specific subregion or cell type of the hippocampus based on current empirical knowledge may be premature. Only feedforward connectivity and one class of input were considered for the purpose of this study, with an abstract model of spiking and plasticity. We nonetheless maintain that this simplified model captures an essential aspect of the translation of spatial information from MEC to HC (see also Savelli and Knierim 2010; Cheng and Frank 2011). The model could apply to DG granule cells (GoodSmith et al. 2017) but this translation might reoccur more than once along the DG-CA3-CA1 trisynaptic loop, where each stage receives direct projections from MEC. Place cells in CA3/1 are additionally influenced by intrahippocampal processing (Bittner et al. 2015; Davoudi and Foster 2019; Fan et al. 2023), which could propagate or refine place tuning (Neher, Azizi, and Cheng 2017). The plasticity-gating role attributed to postsynaptic activity by Eq. 1 is consistent with the artificial induction of place fields in controlled locations via transient juxtacellular or optogenetic stimulation of place cells in DG and CA3/1 (Diamantaki et al. 2016; 2018; McKenzie et al. 2021). These induction protocols do not work on every cell, possibly because a viable grid alignment that can independently sustain firing in the targeted spatial location is not available to all cells. Two additional features of Eq. 1 are the rapid action of plasticity and the ability to heterosynaptically depress inactive or weakly active inputs during postsynaptic activation. Both features are generally compatible with hippocampus physiology (e.g., Remy and Spruston 2007; Levy and Steward 1979; Lynch, Dunwiddie, and Gribkoff 1977). (As our simulations illustrated, however, synaptic reorganization events could unfold over the course of seconds or longer during animal behavior). Particularly relevant to our model are observations that postsynaptic activation need not always take the form of somatic firing in order for place tuning to start emerging intracellularly (D. Lee, Lin, and Lee 2012; Bittner et al. 2017; Cohen, Bolstad, and Lee 2017). It is therefore possible that the model’s principles could apply to dendritic compartments/spikes. Even minor changes in the spatial response of our simulated place cells sometimes correspond to pronounced synaptic changes and reshaping of the landscape of synaptic excitation, which could be tested by monitoring place tuning in real dendrites (Moore et al. 2022). Ultimately, a definitive experimental test of the model would require tracking the evolution of the synaptic efficacies of identified grid inputs during place field formation.

Our theoretical observations characterize place cells as a neural system governed by a tension between plasticity and stability. We showed how the tuning of place cells’ spatial firing patterns and their spontaneous reorganization could be mechanistically indistinguishable. Various physiological correlates of novelty exist in the hippocampal formation (Davis, Jones, and Derrick 2004; Nitz and McNaughton 2004; Frank, Stanley, and Brown 2004; I. Lee, Rao, and Knierim 2004; Barry et al. 2012; Priestley et al. 2022; Hasselmo, Wyble, and Wallenstein 1996), opening the possibility that familiarity with an environment or behavioral context could modulate plasticity and regulate the representational flexibility of the place cell system.

## Model details

### Trajectory data

The head position of male, Long Evans rats was sampled at 30Hz during typical foraging sessions run for place/grid cell experimental studies carried out in the Knierim laboratory at Johns Hopkins University (e.g., (Savelli, Luck, and Knierim 2017) for Plat137). Multiple behavioral sessions were concatenated together to obtain the longer sessions used in our simulations (the same session could occur more than once). The concatenated sessions were always from the same rat and day. To avoid that the input integration and learning dynamics of the simulated cells could be affected by a spurious, instantaneous “teleportation” at the concatenation timepoint, an artificial 10 s pause was inserted between the concatenated sessions. Entries in the position time series corresponding to these pauses were marked as invalid (NaNs). Positions were similarly marked as invalid during pauses naturally performed by the animal, because they are experimentally associated to non-spatial firing in real place cells (rat velocity < 3 cm/s was used as a criterion). During the simulation grid cells were kept from firing spikes at positions marked as invalid during these natural and artificial pauses.

### Grid cells

The triangular grid of each grid cell was defined by the position of one of its vertices (phase), the orientation of the vector linking this vertex with one of its neighbors (orientation), and the intervertex distance (scale). The firing rate of the cell at a given position in space was computed as 20×exp[d²/(0.035×s²)], where *s* is the scale of the grid and *d* is the distance of the position from the closest vertex. The firing rate was used to modulate the instantaneous rate of an inhomogeneous Poisson spike train. Inter-spike intervals shorter than 3 ms were set to 3 ms to create a refractory period.

### Place cells

The membrane dynamics of hippocampal place cells was modeled by

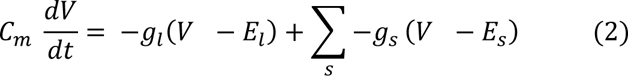

where *V* is the membrane voltage, *C_m_* is the membrane capacitance (2 nF), *g_l_* is the leak conductance (0.2 µS), and *E_l_* is the leak reversal potential (-65 mV). The synaptic contribution to membrane dynamics was modeled with dynamic conductances; for each synapse *s*, *g_s_* is its conductance and *E_s_* is its reversal potential (*E_s_* = 0 mV for all synapses, which are all excitatory in the present simulations). Whenever *V* crossed the firing threshold (-50 mV), a 1-ms spike event was superimposed. For the following refractory period (3 ms), *V* was kept at its resting potential (-70 mV), after which the integration of the voltage dynamics according to Eq. 2 was resumed. The membrane voltage *V* was never allowed to decrease below the lower bound -100 mV or increase above 100 mV (the dynamics of the conductances were not manipulated). Equation 2 was numerically integrated by the exponential Euler method with a 1 ms time step.

In this conductance-based model, the synaptic conductance *g_s_*was 0 in the absence of presynaptic activity. To model synaptic activity, *g_s_* was instantaneously incremented by a quantity *w_s_* every time a presynaptic spike occurred and decayed exponentially to 0 according to the equation *dg_s_/dt = -g_s_/τ_s_*. The time constant τ*_s_*was set to 2 ms, based on the dynamics of AMPA receptors. The value of *w_s_*depended on the previous potentiation or depression of synapse *s* by synaptic plasticity, as described by Eq. 1 (see main text). This variable, therefore, played the role of synaptic weight and was the subject of synaptic modification whenever synaptic plasticity was active in the simulation. The initial value of *w_s_* at all synapses was 4.0×10^-2^ µS. Although starting with identical weights is not biologically realistic, it allows the random set of grid inputs to express their collective untuned pattern of synaptic drive that the model is challenged to turn into place-field-like activity. We refrained from randomizing or constructing a maximally adverse weight distribution for three reasons: (1) to be consistent with the original model; (2) these distributions would not have necessarily been more biologically plausible; and (3) the ability of the model to create new place fields and extinguish old ones midsession described in the present study suggests that the initial weight distribution is not critical to the insights gained from this model. The weights were artificially kept from becoming negative or growing above a saturation limit of 0.1 µS to avoid runaway potentiation. The learning rate *k* in the synaptic rule (Eq. 1) was set to 4 nS/sHz^2^ = 4 nS · s. Synaptic weights were updated every 4 ms according to this rule and saved every 500 ms.

## Analyses details

### Rate maps and place fields detection

Rate maps were computed from trajectory positions and spike trains as in experimental studies (Savelli, Luck, and Knierim 2017). Rat positions were binned in 3 × 3 cm bins to produce an occupancy map of the rat’s dwell time in each bin. The firing rate map was obtained by dividing the count of spikes occurring in each bin by the total occupancy in the same bin. Bins that were never visited were marked (NaN) for exclusion from the map. The rate map was then smoothed by a clipped 2D Gaussian mask with 5 × 5 bins and variance = 2. The values of the mask were dynamically renormalized to account for excluded bins falling within the mask at different steps. If less than five valid bins fell within the mask at any step, the output bin in the smoothed map was marked as unvisited (NaN) and excluded from visualization (white bins in rate maps) and analysis. Place fields were defined as sets of at least 14 contiguous bins with firing rate at least 0.2 Hz and greater than 20% of the highest firing rate of any bin in the rate map (i.e., the peak firing rate of the map).

### Stability Analysis

To compute the PFS (place field stability score) the timestamps of all and only the spikes occurring inside the place field were first organized as a time series. An activity trace for the place field was then calculated as a density estimate of these timepoints via a Gaussian kernel with a 10 s bandwidth. The same process was then applied to the timestamps of the animal positions sampled inside the place field to generate an occupancy trace. In a cell displaying consistent activity in the place field over time, the two traces should wax and wane together. The PFS was thus computed as the normalized dot product of the vector representations of the two traces, which is equivalent to the cosine of the angle between these vectors. Note that, unlike a Pearson correlation, the PFS ranges from 0 to 1 and does not include subtraction of the mean, to prevent meaningless scores in place fields that extend over most of the environment.

To measure total synaptic changes we computed the correlation of synaptic vectors (comprising all weights for all inputs to the cell) at a 10 s lag every 500 ms.

## Code accessibility

Simulations were implemented in C++ and its standard libraries. Data analysis and plotting were implemented in Python using open-source libraries (Numpy, Scipy, Pandas, Matplotlib) made available through the Anaconda distribution (Anaconda, Inc., Austin, TX). Code will be made available via ModelDB and upon request.

## Acknowledgments

This work was started at the Zanvyl Krieger Mind/Brain Institute at the Johns Hopkins University, and it was supported by UTSA startup funds (F.S.) and NIH Grants R01 NS102537 (Noah Cowan, James Knierim, F.S.). I thank Jennifer Siegel, Heekyung Lee, and Chia-Hsuan Wang for sharing animal trajectory data collected in the Knierim Lab. I thank James Knierim for discussion, and Kimberly Christian for discussion and comments on the manuscript.

---

**Supplementary Figure 2_1.**
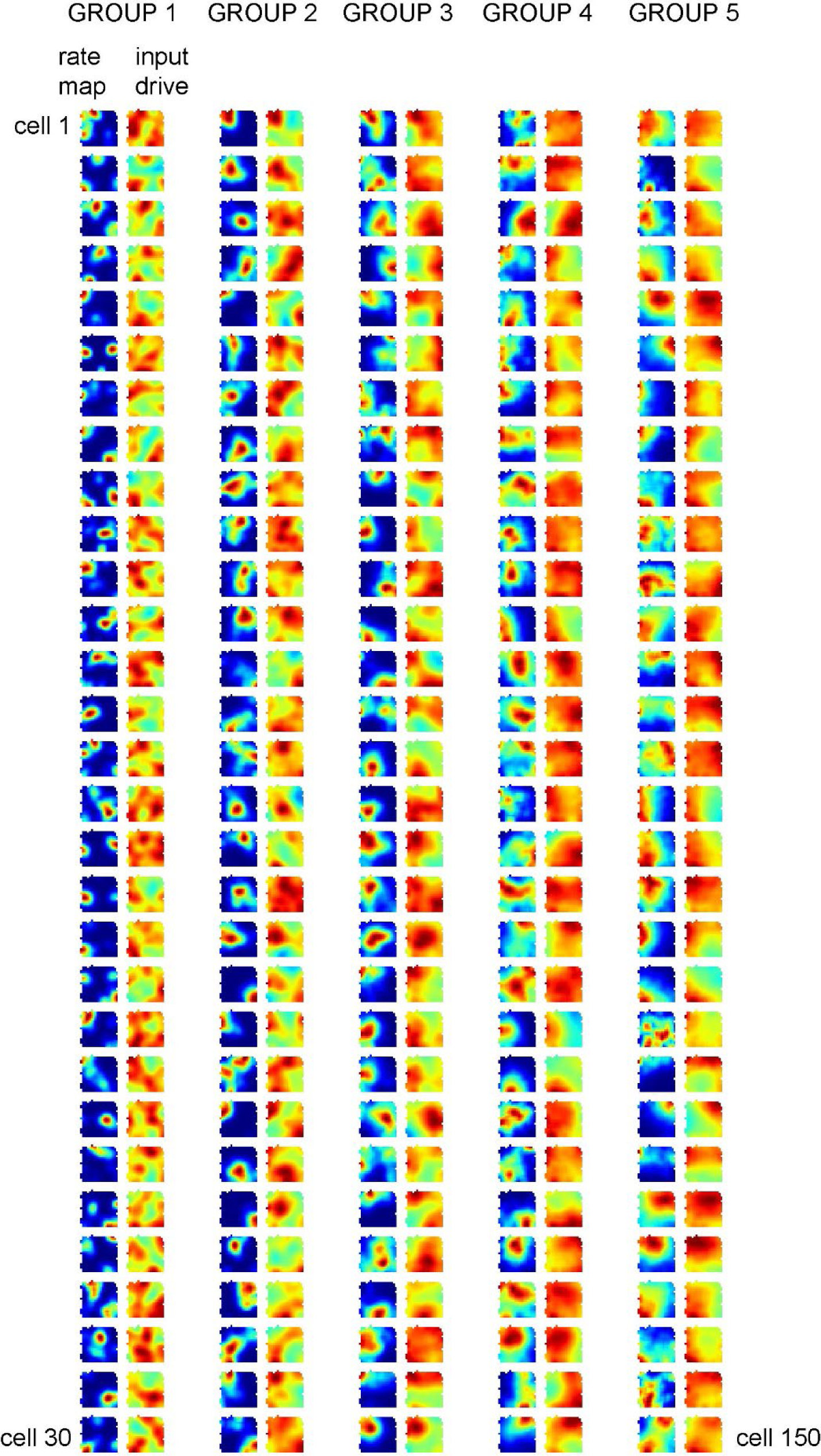
Rate and input drive maps for all place cells co-simulated in Box60.

**Supplementary Figure 2_2.**
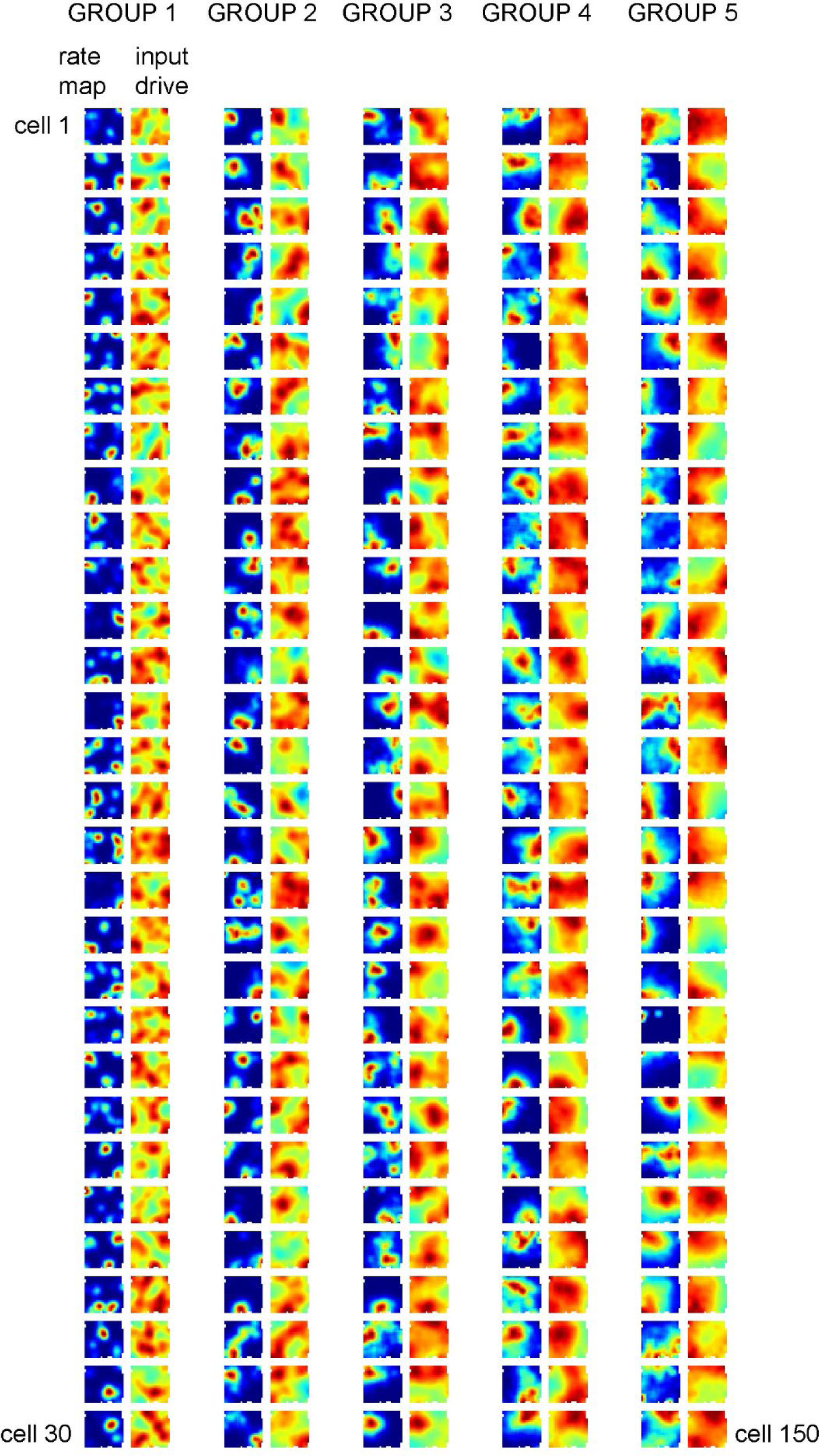
Rate and input drive maps for all place cells co-simulated in Box68.

**Supplementary Figure 2_3.**
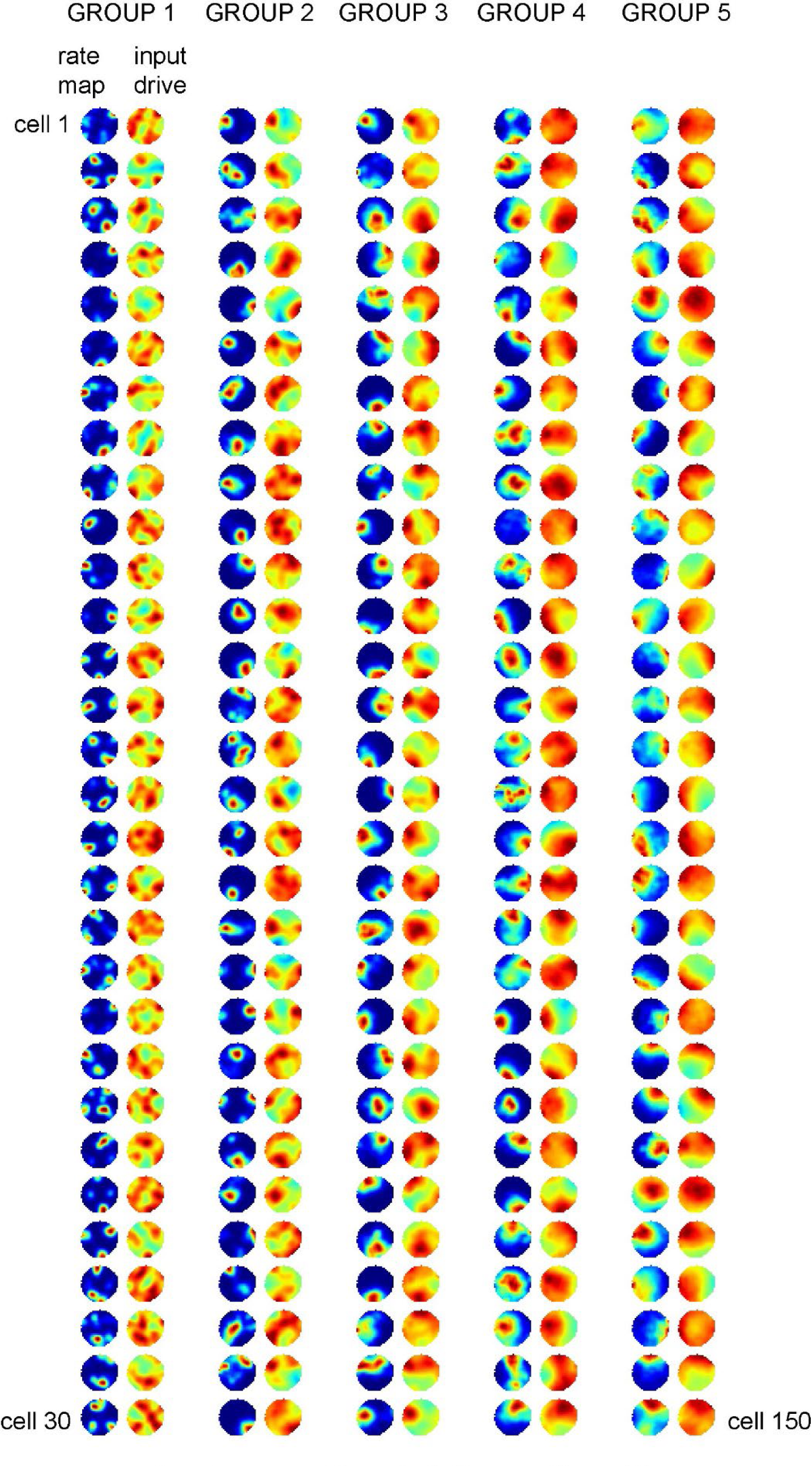
Rate and input drive maps for all place cells co-simulated in Cyl76.

**Supplementary Figure 2_4.**
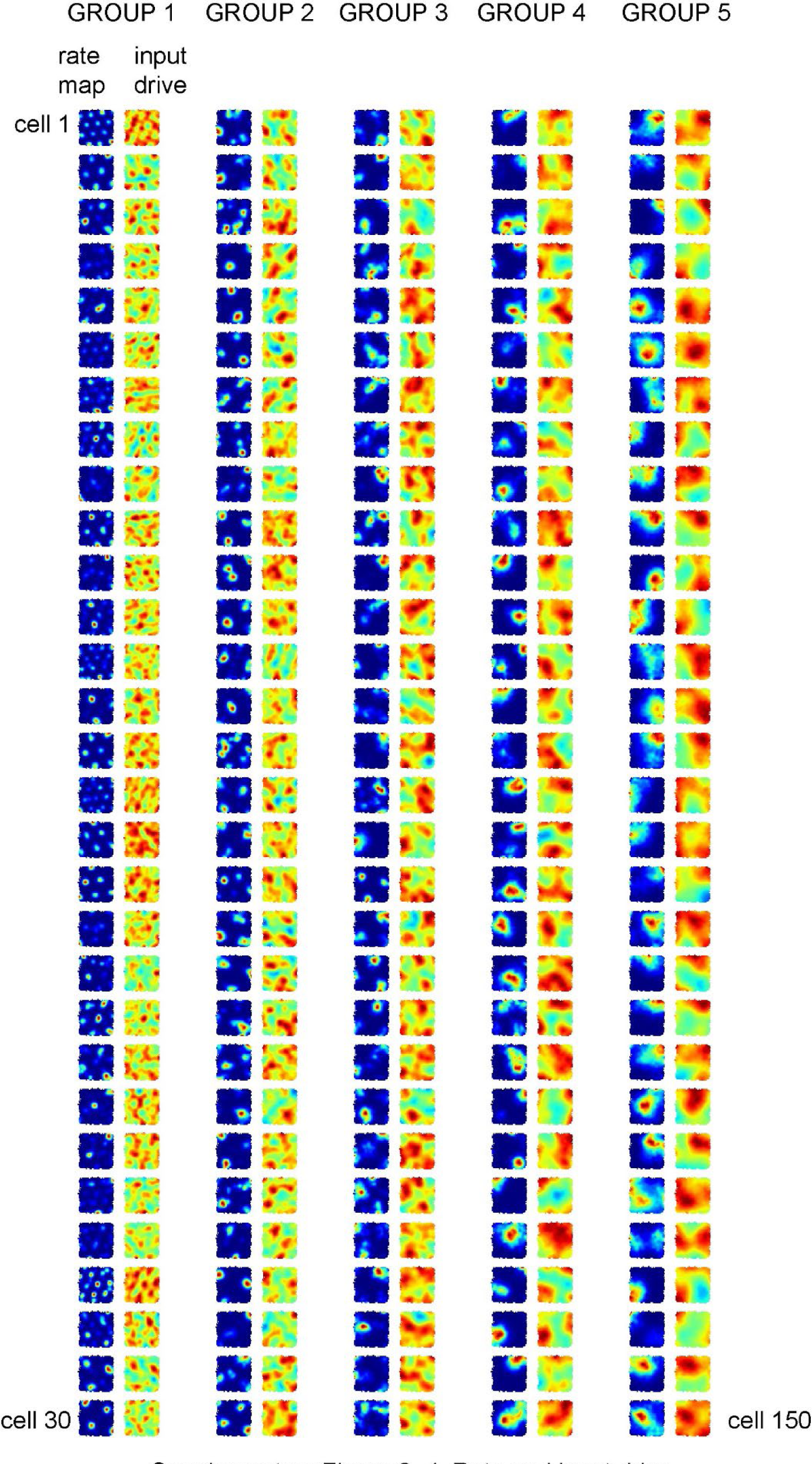
Rate and input drive maps for all place cells co-simulated in Plat100.

**Supplementary Figure 2_5.**
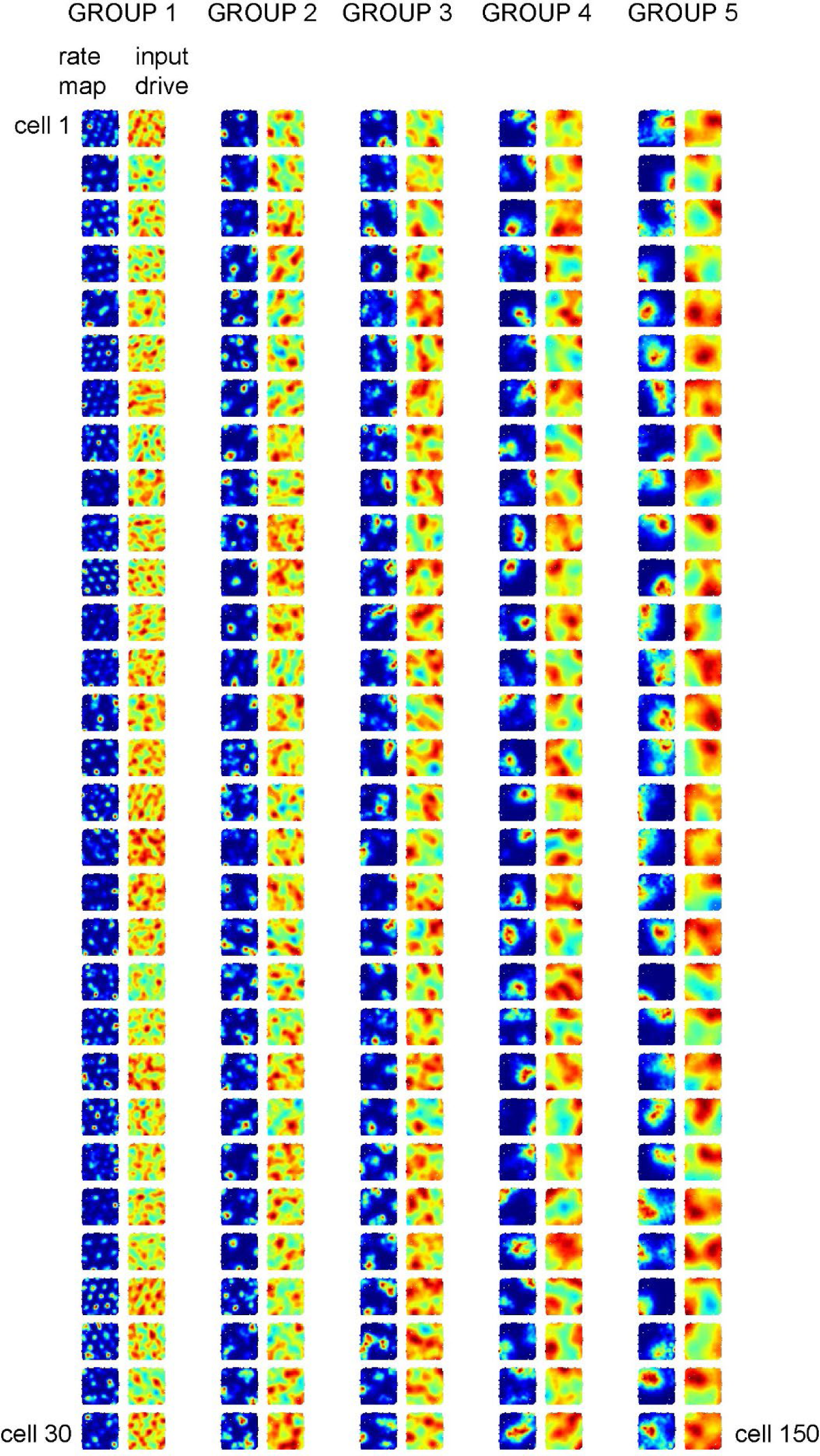
Rate and input drive maps for all place cells co-simulated in Plat137.

